# A flow cytometric approach to identifying the relative abundance and functional capacities of hemocyte subsets in the American cockroach, *Periplaneta americana* (L.)

**DOI:** 10.1101/2024.12.13.628395

**Authors:** Faith J. Boyer-Millander, Aaron T. Martin, Chadwick A. Hamm, Arthur G. Appel, Elizabeth Hiltbold Schwartz

## Abstract

There is growing interest in insects as subjects for comparative immunological studies; however, very little has been done to wholistically characterize insect immune cells using modern techniques such as flow cytometry, and virtually no work of this kind has been done in *Periplaneta americana*. Here, we use an array of general molecular probes including fluorescent lectins, lysosomal indicators, and functional assays to distinguish and characterize insect immune cells (hemocytes) based on cell markers and functions. We have utilized fluorescent tracers of lysosomal content, ROS production, and phagocytosis, as well as microscopic examination of morphology and melanization to distinguish hemocyte types based on these functions. Our findings support the use of lectins as an additional means of separating at least three populations of cockroach immune cells coupled with morphological measurements such as size and complexity. Our results indicate that many functions are enriched in the more granular hemocyte population as these are the cells that undergo phagocytosis and melanization.

## 1. Introduction

Insects, such as *Drosophila*, have historically served as important model systems for immunological studies. The general ease of care and characterization, low cost, lack of regulations, and rapidly reproducing population size are aspects that make insects an appealing system for such studies. Some efforts have been made to investigate the impact of the gut microbiome in development and activity of the innate immune system by generating germ-free *Drosophila*^1,*2*^. However, when considering the complexity of the gut microbiota in conventional animals, *Drosophila* has a relatively simple gut microbiome compared to mammalian systems^3^. The American cockroach (*Periplaneta americana* (L.)) is a global, omnivorous pest with a gut microbiome complexity approaching that of humans^4^. Recently, protocols have been developed to generate axenic *P. americana* making this species an attractive option for investigating the role of the gut microbiome in the development and function of the innate immune system^5,6^. Despite this, little work has been done to wholistically and functionally characterize the hemocyte populations of *P. americana*.

The insect immune system is composed of cells and humoral components interacting within the main body cavity (hemocoel), including fat bodies and immune cells called hemocytes. These cells carry out a variety of innate immune functions including antimicrobial peptide secretion, reactive oxygen species (ROS) production, melanization, and cellular (phagocytosis, encapsulation, nodulation) processes ^7^. Cell populations carrying out these functions have been primarily classified through microscopy; as such, we lack a wholistic and standardized method of cell classification and attribution of immune function to specific cell types in insect systems, as well as an understanding of how their numbers and relative abundances change upon immune challenge.

In mammalian systems, the availability of highly specific reagents such as antibodies allow for a refined classification and investigation of immune cells. Despite being an important and widely used model organism, commercial availability of antibodies to tag immunologically significant targets in *Drosophila* remains limited. This dearth of reagents leads us to seek alternative, non-species-specific methods to investigate insect immune cells. In other systems facing similar challenges, lectins^8^ and ROS indicators^9^ have been used to classify immune cells. Lectin binding activity has been used with hemocytes in recent years; however, there has not been a wide-spread effort to use these reagents as a means of defining or sorting hemocyte populations^10-12^. Additionally, while infrequently used^12-14^, flow cytometry has largely been underutilized as a method of investigating insect immune cells. In this study, we employ the use of flow cytometry and cell sorting in tandem with a variety of general reagents (lectins, lysosomal indicators, ROS indicators) and functional assays (phagocytosis, melanization) to distinguish hemocyte populations based on specific cell characteristics and functions. Our findings provide a necessary foundation for the characterization of the *P. americana* immune system for future immunological studies, including the impact of the gut microbiome in insect immunity. Additionally, our methods establish a means of investigation of hemocytes that can be more broadly applied in other insect systems.

## 2. Materials and Methods

### 2.1 Animals

*Periplaneta americana* used in this study were maintained as described previously by Smith et al. 2022^15^. Briefly, *P. americana* were housed in 32-gallon Rubbermaid trash bins as mixed sex and age colonies. These colonies were maintained at 27 ± 2°C on a 12:12 L:D photoperiod. Rolled corrugated cardboard housing was provided as harborage, and Purina Laboratory Diet 5001 rat chow and water provided *ad libitum*. Males were individually captured from the colony by hand.

### 2.2 Hemolymph collection

Animals were anesthetized on ice for 5-7 minutes prior to hemolymph collection. Hemolymph was collected by cutting the coxal-trochanteral joint of the metathoracic legs and pipetted into sterile-filtered anticoagulant buffer (69 mM potassium chloride, 27 mM sodium chloride, 2 mM sodium bicarbonate, 30 mM trisodium citrate dihydrate, 26 mM citric acid, 10 mM edetic acid)^16^. Samples were pooled or collected individually as indicated into sterile 1.5mL microcentrifuge tubes.

### 2.3 Flow cytometric analysis and cell sorting

Whole hemolymph was strained into 5 mL polystyrene round-bottom tubes through a cell strainer cap (Corning 352235) before use in flow cytometry. Hemocytes were analyzed on a MACSQuant Analyzer 10. Cell sorting was carried out on a CytoFLEX SRT at the AUCVM Flow Cytometry Facility.

Sorted hemocyte samples were collected in 5 mL polypropylene tubes (Sarstedt 55526006) containing 1mL of serum-free HyClone CCM3 Insect Media (Cytiva SH30065.02). Doublets and debris were excluded from analysis (**Supplemental Figure 1)**. Analysis of flow cytometric data was conducted using FlowJo 10.

### 2.4 Staining with fluorescent lectins, DCFDA, and lysosomal indicator

#### Lectin staining

Hemocytes were collected into 100 µL anticoagulant buffer and insect media was added up to 1 mL. A panel of fluorescent lectins (ConA, WGA, UEA I, GSL I, and LEA) were used at a final working concentration designated by the manufacturer protocol (**Table 1)** (Invitrogen). Each sample was incubated at 30°C for 10 minutes and washed twice at 500 x g for 5 minutes in insect media before being resuspended in anticoagulant buffer. Cells were passed through a cell strainer cap prior to analysis via flow cytometry or cell sorting.

**Table 1.** Lectins used to evaluate lectin binding capabilities of hemocytes. Table includes the full lectin name, the lectin abbreviation used throughout this paper, the concentration, and the catalog number from the manufacturer.

| Lectin full name | Abbreviation | Concentration (ug/mL) | Cat. no (Invitrogen) |
| --- | --- | --- | --- |
| Concanavalin A | ConA | 100 | C56127 |
| Wheat germ agglutinin | WGA | 10 | W56132 |
| *Ulex europaeis* agglutinin I | UEA I | 20 | L32476 |
| *Griffonia simplicifolia* lectin I | GSL I | 10 | L32473 |
| *Lycopersicon esculentum* agglutinin | LEA or LEL | 10 | L32472 |

#### DCFDA staining

To evaluate the production of reactive oxygen species in hemocytes, cells were collected as described above and stained with CM-H_2_DCFDA (Thermo C6827), a general oxidative stress indicator, according to manufacturer protocol. Briefly, hemocytes were collected as described above and incubated with DCFDA at a working concentration of 10 µM for 20 minutes at 30°C. Each sample was washed at 500 x g for 5 minutes and resuspended in an anticoagulant buffer prior to analysis through flow cytometry.

#### LysoTracker™ staining

Lysosomal content of hemocyte populations was investigated using LysoTracker™ (Invitrogen L7528) staining according to manufacturer protocol. Briefly, hemocytes were collected as described above and incubated with LysoTracker™ at a working concentration of 75 nM for 2 hours at 30°C. Hemocytes were washed at 500 x g for 5 minutes, resuspended in anticoagulant buffer, and analyzed via flow cytometry.

### 2.5 Fluorescent staining and microscopy

Sorted hemocytes were incubated with Phalloidin (F-actin, AAT Bioquest 23153) for 30 minutes at 30°C in a sterile microcentrifuge tube and washed 2x with insect media. The cell pellet was resuspended in ProLong^TM^ Gold antifade mounting media with DAPI (Invitrogen P36935) and placed on a cover slip prior to imaging. Microscopy and imaging were carried out using a Zeiss Axio Observer ApoTome.2.

### 2.6 Melanization and microscopy

To stain prophenoloxidase-containing cells in whole hemolymph, hemolymph was collected from ten adult American cockroaches and pooled in one mL of anticoagulant. Following this, 40 µl of pooled whole hemolymph samples with (experimental group) or without (control group) L-DOPA or ethanol were placed in one well within a 96-well plate and incubated at 30°C for two hours across three experiments. For experimental groups, this mixture contained 100 µl of a 2 mg/mL 3,4-Dihydroxy-L-phenylalanine 98+% (L-DOPA) solution (Thermo Scientific, A11311.22) suspended in Dulbecco’s Phosphate Buffered Saline (DPBS, Fisher Scientific, BW17-512F), 60 µl of 70% denatured ethanol, and 40 µl of pooled whole hemolymph for a total volume of 200 µl. The rationale for the addition of L-DOPA and ethanol is their significance in facilitating hemocyte staining. The melanization reaction relies on the presence of the L-DOPA substrate to proceed^17^ and ethanol has been described to spontaneously activate the prophenoloxidase (PPO) enzyme^18^, an integral protein in the enzymatic cascade that synthesizes in cytotoxic melanin. In contrast, control groups contained 160 µl of DPBS and 40 µl of pooled whole hemolymph. These samples were observed via brightfield microscopy, and five fields were recorded.

Following cell sorting based on the LEA lectin staining, each separated population was seeded in triplicates in a 96-well plate at a density of 50,000 cells in 200 µL of insect media. After overnight incubation, brightfield microscopy of the hemocyte populations was used to establish a control level of hemocyte darkening for each sample. Following microscopic examination of the control samples, media from all wells was aspirated and a mixture of ethanol and L-DOPA was added to induce hemocyte staining. The wells of this hemocyte staining assay contained 100 µl of L-DOPA solution, 60 µl of 70% denatured ethanol, and 40 µl of insect media for a total volume of 200 µl. Following the addition of the reagent cocktail, all samples were incubated at 30°C for two hours and then brightfield microscopy images were taken to record phenotypic changes across all hemocyte populations. The percentage of individual stained cells within each hemocyte group was calculated by applying a grid to each microscopy image (ten images per replicate), randomizing the squares counted via a random number generator, and counting both PPO-containing cells and total cells within the image across three experiments.

### 2.7 Phagocytosis

The assay used to measure phagocytosis was modified from a previous study^19^. Briefly, hemolymph was collected and pooled from a sample of five to ten cockroaches. Hemocytes were then counted and checked for viability using trypan blue exclusion dye (Invitrogen K940). Approximately 85,000 cells were then seeded into a 96-well plate (VWR 734-2327) with 190 µL of CCM3 insect media in each well. Then, 10 µL of pHrodo™ Bioparticles® (Invitrogen P35367), *Staphylococcus aureus* bacteria conjugated with a pH sensitive fluorophore, were added to each experimental well. The cells were then incubated for 2, 4, 8, 16, or 24 hours with the bioparticles at 25°C. To collect samples at each timepoint, the cells are lifted from the plate using cold PBS, stained with LEA for 10 minutes, and centrifuged at 500 x g for 5 minutes. The supernatant was removed, and the pellet was resuspended in 500 µL of PBS to be acquired on the flow cytometer

## 3. Results

### 3.1 Relative abundance of hemocyte subsets measured by flow cytometry

To determine if distinct hemocyte subsets could be visualized by examining the forward scatter (cellular size) and side scatter (intracellular complexity) of whole hemolymph, hemocytes were collected from 19 individual *P. americana* into anticoagulant buffer, filtered, and immediately analyzed by flow cytometry. Doublets and debris were excluded during analysis, and hemocytes were plotted on a dot plot comparing cell size and complexity **(Figure 1A)**. Generally, five populations of hemocytes (indicated by gates on Figure 1A) could be discerned based on these characteristics. The dot plots in Figure 1A depict the cellular size and complexity patterns of four representative individuals. As demonstrated here, the relative abundance of each population was highly variable between individuals **(Figure 1A)**. With these groupings, we observed that the “small” hemocyte population was the most abundant, but also the most variable. “Large” cells were also highly abundant with more consistency, followed by “granular,” “large/complex,” and “low complexity” groups **(Figure 1B)**. Due to the high variation between individuals, size and complexity were determined to be too inconsistent as a basis for cell sorting. Thus, we sought other methods and reagents to obtain consistently labeled populations.

**Figure 1.**
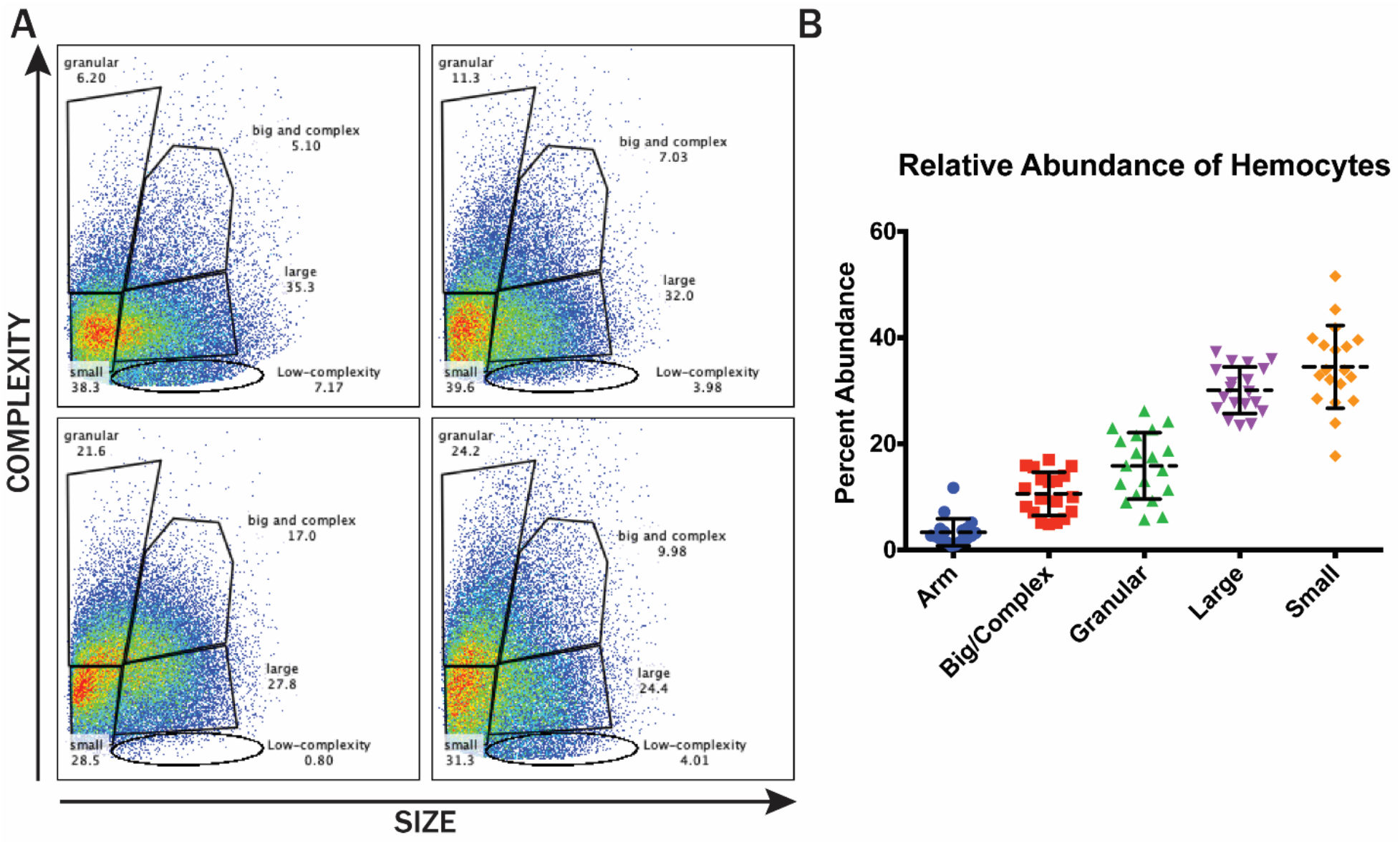
Flow cytometric analysis of *P. americana* hemocytes based on cell size and complexity. **(A)** Four representative plots of individual *P. americana* hemocytes displaying variability between individuals. Generally, five populations were regularly observed (small, large, granular, big/complex, low complexity). **(B)** Relative abundance of hemocyte populations observed across individual *P. americana* (N=19). “Small” cells were the most abundant, but variable, and low complexity was a comparatively rare population.

We next tested whether lectin binding would serve to distinguish hemocyte populations. We employed a panel of lectins with different binding specificities (WGA, LEA, ConA, GSL I, and UEA I) to distinguish *P. americana* hemocyte populations. Applying this panel, we found that hemocytes bound WGA (binds both 1,3-N-acetylglucosamine and N-acetylneuraminic acid), LEA (binds 1,3-N-acetylglucosamine), and ConA, with the greatest separation of fluorescent peaks observed with LEA (*Lycopersicon esculentum* agglutinin) and WGA (Wheat-germ agglutinin). There was no evidence of any hemocyte population binding GSL I or UEA I **(Supplementary Fiigure 2)**. Compared to the unstained control **(Figure 2A)**, we observed that WGA binding distinguished two populations of *P. americana* hemocytes, one WGA positive and one WGA negative group **(Figure 2B)**. In contrast, LEA binding distinguished three hemocyte populations, LEA-low, medium, and high-binding groups **(Figure 2C)**. When both lectins were used to stain the cells, the pattern of staining was generally either double negative (-/-) or double positive (+/+) **(Figure 2D)**. When we examined these subsets based on size and complexity, we found that the -/- group was a less complex group, while the +/+ cells were primarily granular cells **(Figure 2E)**. Based on its ability to separate three populations, we then chose to utilize LEA binding alone against cell complexity to obtain three consistent cell subsets **(Figure 2F)**. Relative abundance of three grouped based on LEA-binding capabilities were relatively consistent across individuals, with “LEA-high” being the most abundant **(Figure 2G)**. Cell sorting based on the same parameters revealed enrichments of specific hemocyte morphologies in each group as represented in micrographs, where LEA-low non-complex were small with very little cytoplasm, LEA-low mid complex had a distinct prolate spheroid shape, and LEA-high cells were generally larger with more cytoplasm **(Figure 2H)**. Thus, we determined that lectin staining in combination with cellular complexity enabled us to distinguish three populations of cells that proved to have distinct morphologies upon microscopic evaluation.

**Figure 2.**
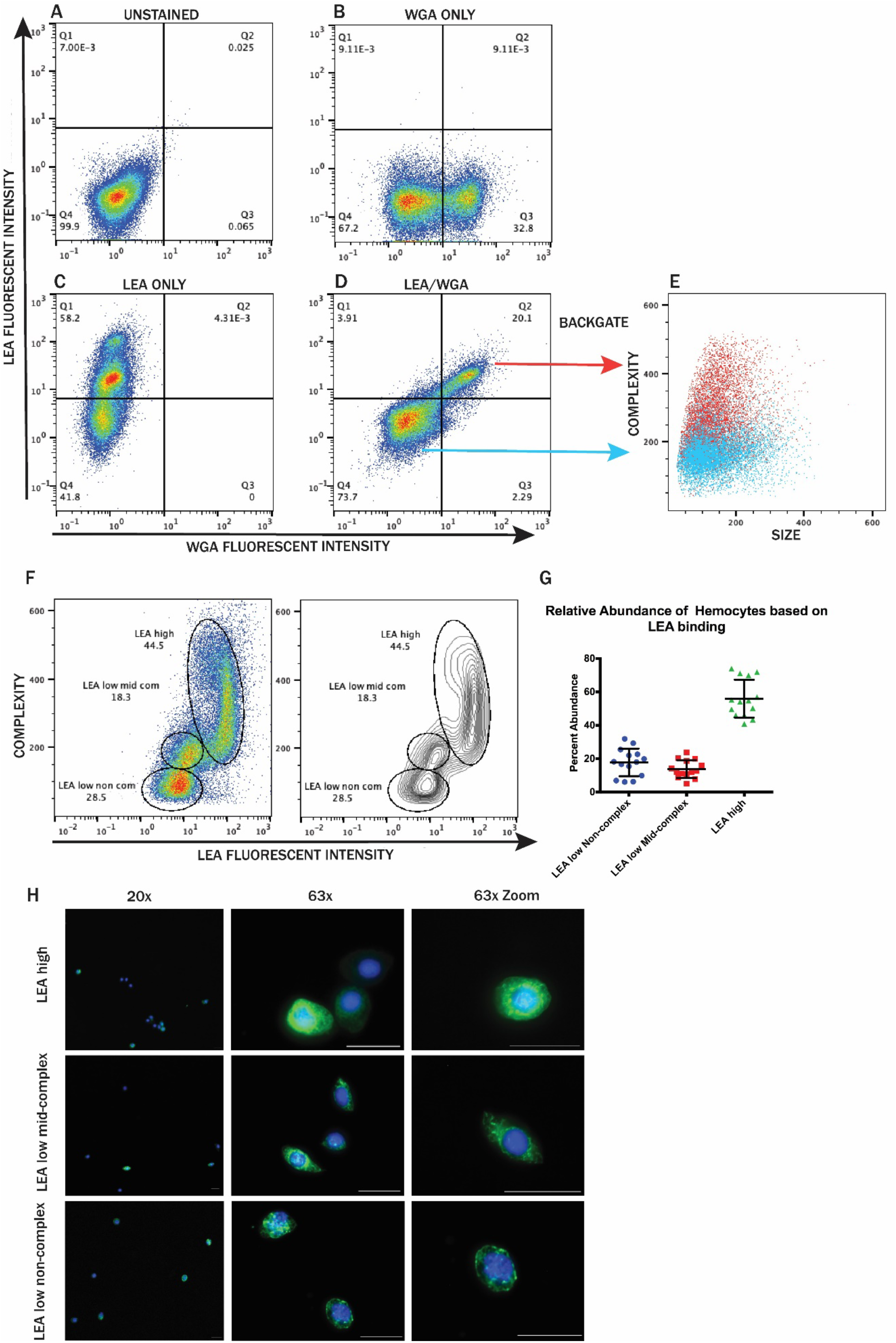
Flow cytometric analysis of *P. americana* hemocytes based on lectin binding capabilities. **(A)** Unstained, **(B)** LEA (*Lycopersicon esculentum* agglutinin) stained, **(C)** WGA (Wheat-germ agglutinin) stained, and **(D)** double stained hemocyte populations. Two populations were distinguished by WGA binding, while three were revealed by LEA binding. **(E)** +/+ cells are more granular, while -/- are comparatively non-complex. **(F)** Representative dot and contour plots based on LEA binding. Three clusters were observed (LEA-low non-complex, LEA-low mid-complex, LEA-high). **(G)** Compiled data from 14 individuals depicting relative abundance of hemocytes based on LEA binding across individual *P. americana*. **(H)** Microscopy of sorted hemocyte populations based on LEA binding. Micrographs represent enriched hemocyte populations in each group. Scale bar is 20 µm.

### 3.2. Lysosomal content and ROS production

To begin to measure functional capacities of hemocyte groups using flow cytometry, we used two fluorescent tracers, LysoTracker™ to quantitate lysosomal content and DCFDA to detect ROS production. Lysosomal content was evaluated across individual cockroaches, and the LysoTracker™ stain generally tagged a population of hemocytes with greater fluorescent intensity, indicating the presence of low pH compartments **(Figure 3A)**. However, the percentage of cells staining brightly varied between individuals, with the lyso-positive population comprising between 14-33% of the total population **(Figure 3D)**. The LysoTracker™ positive and negative groups were then examined for their relative size and complexity. We found that the LysoTracker™ positive peak contained primarily granular cells **(Figure 3B)**, while the LysoTracker™ negative cells were typically non-complex **(Figure 3C)**. Thus, the cells with the greatest lysosomal content were also those with the highest intracellular complexity, as might be predicted.

**Figure 3.**
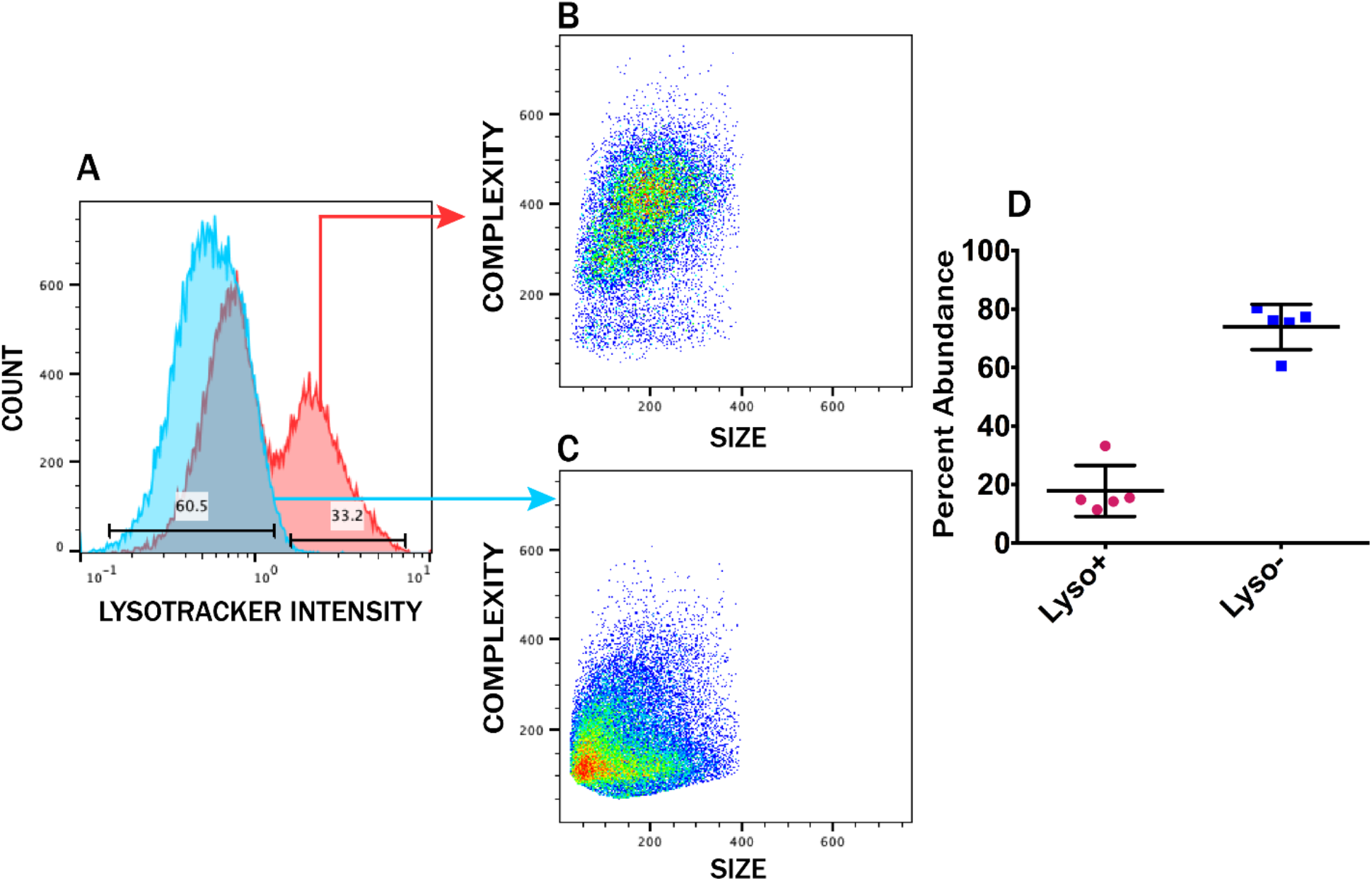
Flow cytometric analysis of lysosomal content. **(A)** Representative graph of unstained (blue) or LysoTracker™ stained (red) samples. When evaluated for cell size and intracellular complexity, the **(B)** lyso-positive peak was primarily granular cells, while the **(C)** negative peak was primarily non-complex cells. **(D)** Relative abundance of LysoTracker™ positive and negative cells (N=5). The majority of hemocytes are lyso-negative, with the lyso-positive peak comprising between 14-33% of the total population.

To evaluate hemocytes based on cellular complexity and ROS production, we stained hemocytes with DCFDA and monitored their fluorescence through flow cytometry. Three populations were distinguishable: ROS low, ROS high granular, and ROS high non-complex **(Figure 4A)**. The ROS low population was the most abundant, making up approximately 50 percent of the population **(Figure 4B)**. ROS high granular and ROS high non-complex were approximately equal in abundance at around 20 percent; however, the granular population was slightly more variable, and this group sometimes had a brighter fluorescent intensity than the non-complex group **(Figure 4C)**. Connecting LEA binding capabilities to ROS production, we found that double staining with LEA and DCFDA revealed two populations of ROS-producing cells: an LEA high and an LEA low group **(Figure 4D)**. When evaluating these cells based on cell size and intracellular complexity, we found that LEA high ROS high cells were more granular, while LEA low ROS high cells were relatively non-complex **(Figure 4E)**. We therefore concluded that two different hemocyte subsets are capable of producing high levels of ROS, and these groups are additionally distinct in both LEA binding capacities and cellular size and complexity.

**Figure 4.**
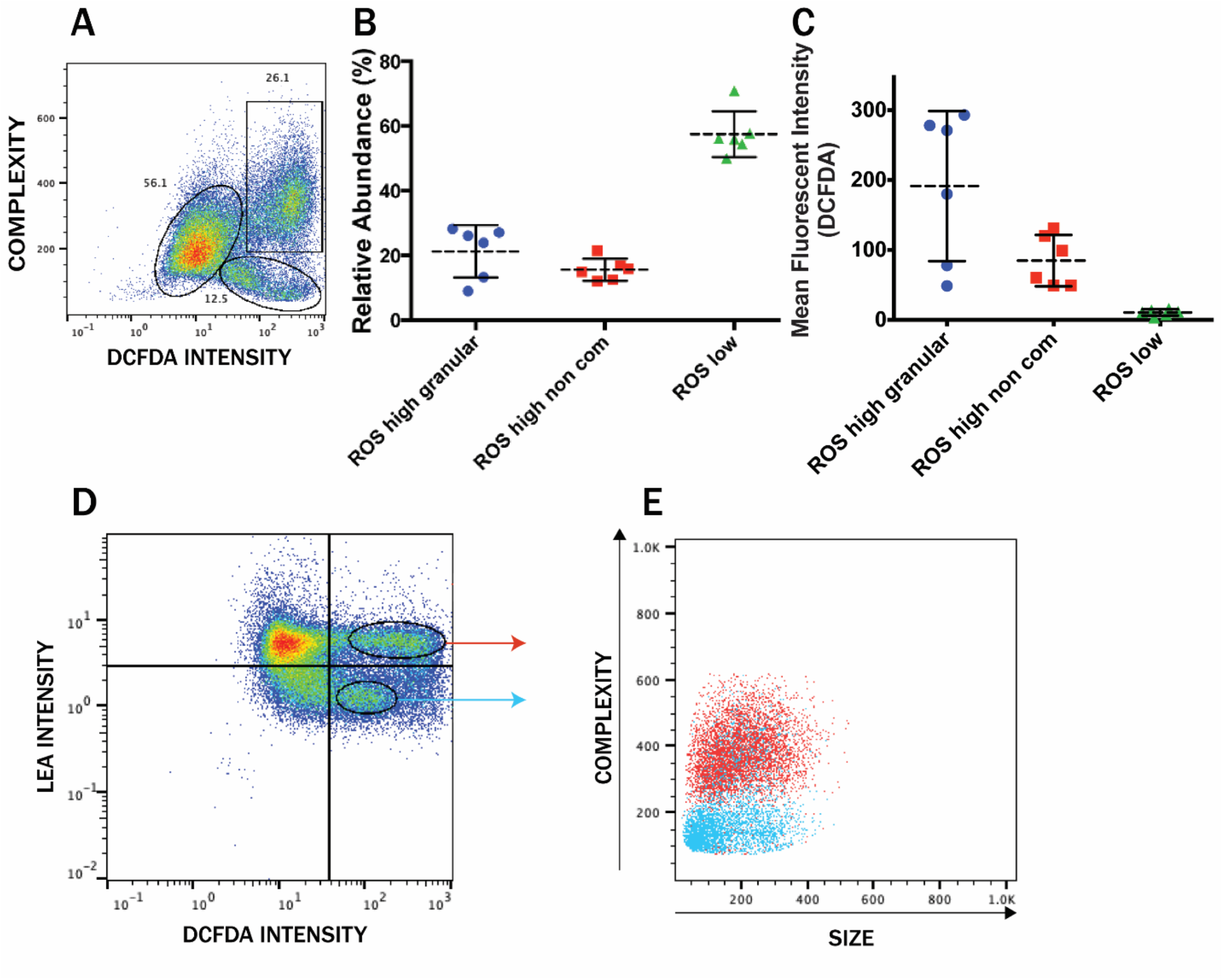
Flow cytometric analysis of ROS production. **(A)** Representative dot plot of hemocyte populations based on ROS production and intracellular complexity immediately after cell collection and staining. There are three populations: ROS high granular, ROS high non-complex, and ROS low. **(B)** Relative abundance of hemocyte populations based on intracellular granularity and ROS production intracellular granularity and ROS production. **(C)** Mean fluorescent intensity (DCFDA) of hemocyte populations distinguished by ROS production. ROS high granular is the most variable group. **(D)** Hemocyte populations distinguished through double staining with DCFDA and LEA. Two populations of ROS-producing cells had differential LEA binding capacities. **(E)** Hemocyte populations distinguished by ROS production and LEA binding affinity evaluated based on size and complexity. The LEA-high ROS-producing cells are generally more granular, while the LEA-low ROS-producing cells are relatively non-complex.

### 3.3 Hemolymphatic melanization and lectin affinity

Our next goal was to identify the cells involved in melanization among *P. americana* hemocytes. The primary enzyme driving the synthesis of cytotoxic melanin is phenoloxidase (PO). Phenoloxidases must first be activated from their zymogen form, prophenoloxidases (PPO) before function. Experimentally, this can be achieved through the addition of a low concentration of ethanol (EtOH). Further, the melanization reaction is mediated by phenoloxidase (PO) acting on L-3,4-dihydroxyphenylalanine, also known as L-DOPA. Used together these reagents serve to activate prophenoloxidase (EtOH) and as a substrate for the reaction (L-DOPA), identifying prophenoloxidase content within a cell and therefore its ability to mediate melanization. Thus, darkening of a cell after exposure to these reagents is directly correlated to prophenoloxidase expression by that cell. To demonstrate this first in whole hemolymph, two whole hemolymph sample groups were collected into anticoagulant buffer and incubated with or without ethanol and L-DOPA for two hours. These samples were then observed with brightfield microscopy. Within the control group where staining was not induced, little to no hemocyte darkening was observed in comparison to the background **(Figure 5A)**. In contrast, the induced group demonstrated a high number of stained cells **(Figure 5B)**. Thus, L-DOPA and ethanol induced hemocyte staining effectively, yet this staining was observed in cells with several different morphologies. To better identify the hemocytes with this capacity, we next utilized a cell sorting approach.

**Figure 5.**
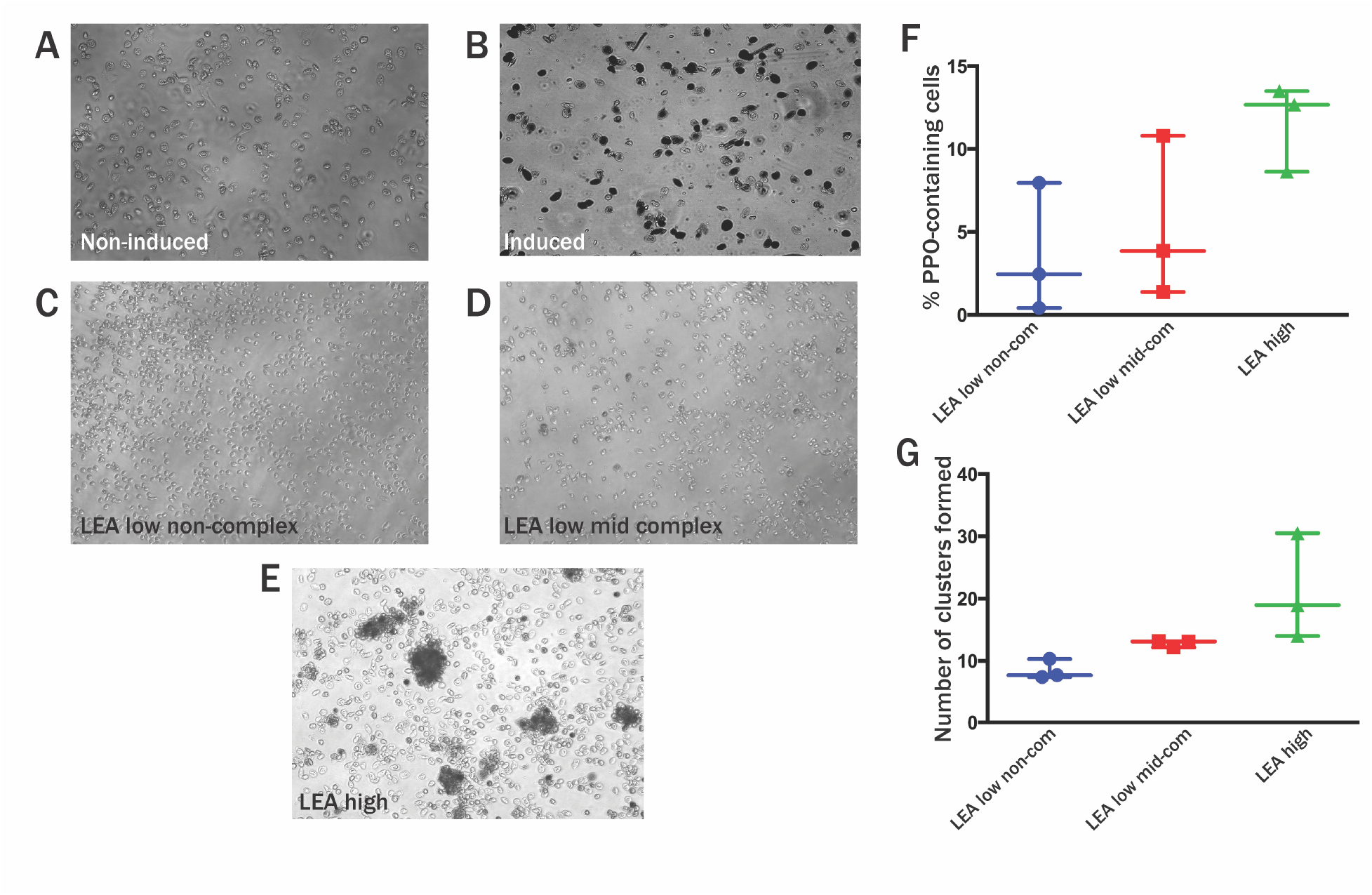
Representative images and quantities of PPO-expression and clustering of hemocyte subsets isolated by cell sorting. (**A and B**) Representative images of a non-induced (no ethanol or L-DOPA) whole hemolymph population (**A**) and an induced (with L-DOPA and ethanol) whole hemolymph population (**B**). (**C, D, and E**) Identification of PPO-expressing hemocyte populations enriched via cell sorting based on binding affinity to LEA. **(F)** Quantitation of the percentage of individual PPO-expressing cells across hemocyte subsets (Blue: LEA-low, non-complex, Red: LEA-low, mid-complex, Green: LEA-high). (**G**) Quantitation of hemocyte clusters formed across hemocyte subsets. Clusters are defined as the aggregation of four or more hemocytes (Blue: LEA-low, non-complex, Red: LEA-low, mid-complex, Green: LEA-high).

To identify cells expressing prophenoloxidase, we sorted hemocyte subsets based on LEA binding and intracellular complexity (using the gating strategy of figure 2F). Three subsets were obtained (LEA low, non-complex, LEA low, mid-complex, and LEA high) via cell sorting. After incubation with L-DOPA and ethanol, these subsets were evaluated for the number of dark stained cells present by microscopy. In the LEA low, non-complex subset, there were little to no dark stained cells **(Figure 5C)**. Among the LEA low, mid-complexity subset there were a few dark stained cells **(Figure 5D)**, and the LEA high subset contained the highest number of stained cells **(Figure 5E)**. We then evaluated the percentage of individual PPO-expressing cells present within the hemocyte subsets by counting the number of dark cells across ten microscopic fields for each subset across three experiments. A grid was applied to each microscopy image that separated into 20 squares. Ten squares of each grid were selected using a random number generator and the number of stained vs. total cells were calculated. These numbers were then averaged with each data point representing one experiment **(Figure 5F)**. Microscopic examination of each hemocyte subset revealed that the LEA-high population had the highest percentage of dark single cells (11.7%) while LEA-low non-complex had the least (2.94%) **(Figure 5F)**. The LEA low, mid-complexity subset showed an intermediate level of stained cells, with these hemocytes comprising 5.73% of the total population **(Figure 5F)**.

We also observed that among some of the hemocyte subsets, there was a high degree of cell clustering upon culture and staining. To quantitate this activity, we averaged the number of clusters observed in ten fields on non-grided images per experiment across three experiments. Each data point is representative of one experiment **(Figure 5G)**. Hemocyte clustering was highest in the LEA-high subset, the LEA-low mid-complex subset had a low amount of clustering, and there was little clustering observed in the LEA-low non-complex population. **(Figure 5G)**. The LEA-high hemocyte subset had a unique activity in the generation of large, stained aggregates **(Figure 5E)**. Thus, the LEA-high subset was enriched in melanizing activities, PPO-expression and cluster formation, that is reminiscent of nodulation. Conversely, hemocyte populations characterized by a low affinity to LEA demonstrated little to none of these melanization capacities. Next, we sought to characterize the hemocyte subsets responsible for another immunological function, phagocytosis.

### 3.4 Phagocytosis

To determine which cell populations within the hemolymph were capable of phagocytosis, we used *Staphylococcus aureus* pHrodo^TM^ BioParticles® to identify those hemocytes that take up and acidify cargo. These particles become progressively more fluorescent as the pH drops, with maximal fluorescence around pH 4. The cells that were able to take up the particles had clearly visible fluorescence within the cells, while exogenous particles outside of the cells were nonfluorescent **(Figure 6A)**. There were noticeable filopodia and lamellipodia-like sheets coming from the cells that had taken up particles. Thus, this reagent enabled us to distinguish cells that contained acidified particles vs. those that did not **(Figure 6A)**. Using flow cytometry, we observed that when staining with LEA and pHrodo^TM^ over a 24-hour period, we observed approximately 15% of the total cell population become pHrodo^TM^ positive **(Figure 6B)**. The percentage of cells with pHrodo^TM^ fluorescence increased progressively over the 24-hour time course, with the largest increase occurring between 16 and 24 hours **(Figure 6C)**. Both the number of cells and mean fluorescent intensity (MFI) increased over the duration of this time course. These phagocytes may have taken up and acidified multiple particles, as the normalized MFI increased to over two times greater than its unstained counterpart compared to other earlier time points (**Figure 6D)**. Flow cytometry of the pHrodo^TM^ with a co-stain of LEA revealed a pattern of two populations, an LEA high group and an LEA low group **(Figure 6E)**. We then wanted to determine if these groups had distinct capacities in uptake or kinetics of acidification. However, due to the variability between biological replicates, we observed no significant differences between the subsets in these capacities **(Figure 6F)**. Yet, the clear separation of phagocytosis-positive cells based on LEA binding intensity led us to question whether these were two separate cell types. When these LEA high and low subsets were further evaluated based on size and complexity, the two were easily distinguishable. The high LEA group contained larger and more complex cells **(Figure 6G)**; while the LEA low group was smaller and less complex **(Figure 6H)**. This could indicate that there are two separate cell types capable of phagocytosis. Further studies will be required to further define the functional identities of these two cell subsets.

**Figure 6.**
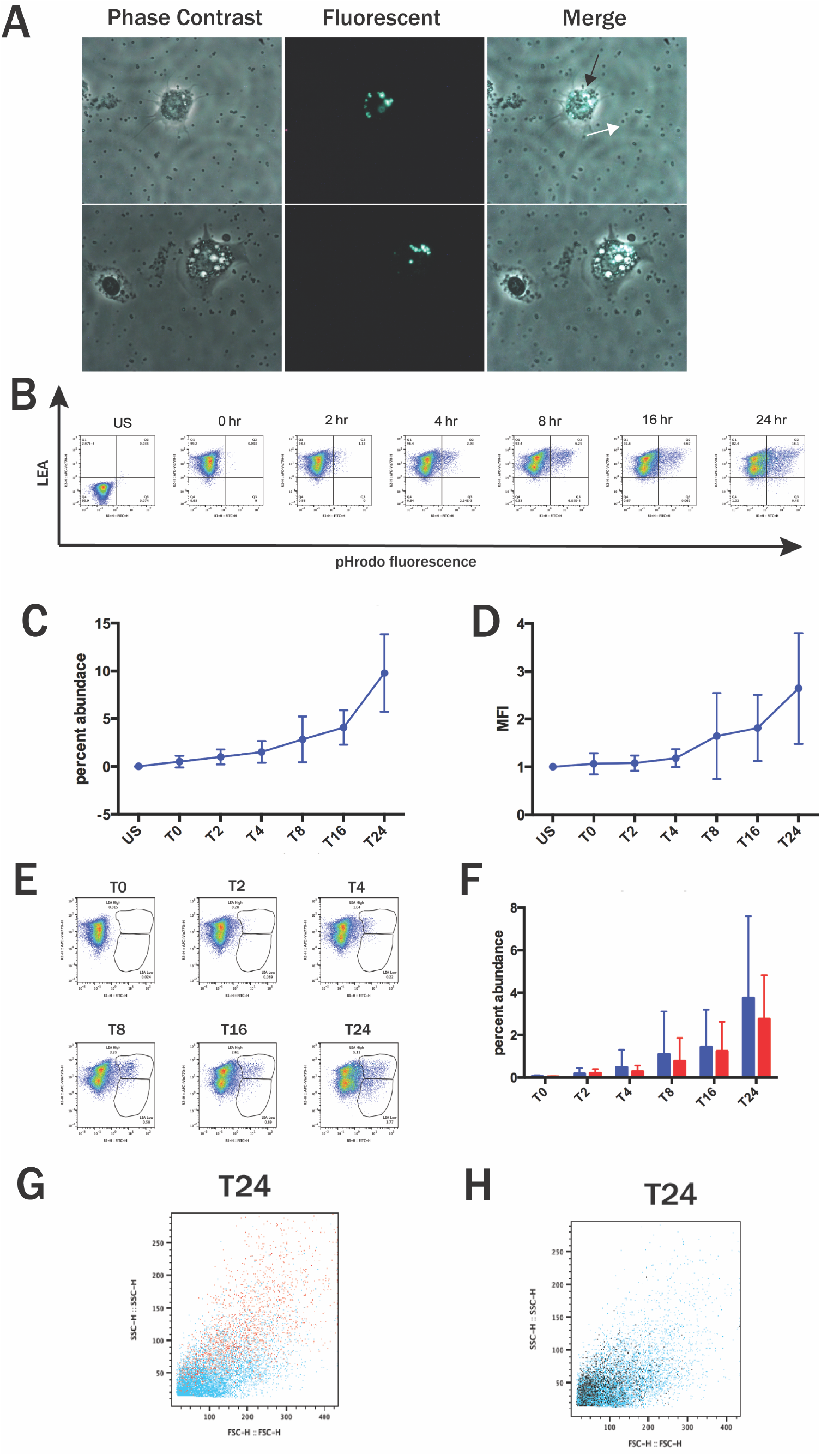
Identification of phagocytic cells using pHrodo^TM^ and LEA. **(A)** Microscopy of phagocytic cells after incubation with pHrodo^TM^, exogenous particles (white arrow) and phagocytized particles (black arrow) **(B)** Representative dot plots of a time course of phagocytosis using pHrodo^TM^ (x-axis) and LEA (y-axis) over a 24-hour period with timepoints at 0, 2, 4, 8, 16, and 24 hours. **(C)** Quantitation of the percent abundance of LEA+/pHrodo+ cells over time. **(D)** Fold change quantitation of the mean fluorescence intensity (MFI) of the LEA+/pHrodo+ cells normalized against unstained control (US). **(E)** Representative dot plots of the separation of the pHrodo+ cells into LEA high and LEA low gates over a 24-hour time course. **(F)** Quantitation of the percent of cells within the LEA high and LEA low gates. **(G)** Representative overlay of ungated hemocytes (blue) and LEA high gates (red). **(H)** Representative overlay of ungated hemocytes (blue) and LEA low gates (black). N=6

## 4. Discussion

While previous studies have identified hemocyte types or subsets within the hemolymph of *P. americana* based on morphology, the relative abundance of these subsets has not been explored. Likewise, the functional capacities of specific cell subsets remain incompletely defined. The application of flow cytometry techniques to these questions has the potential to more wholistically illuminate the composition and functional abilities of these cells. Previous microscopy-based studies in *P. americana* have reported the presence of prohemocytes, granulocytes, plasmatocytes, oenocytoids, spherulocytes, and adipohemocytes^20,21^. Our initial flow cytometric analyses employing size and complexity as defining cell characteristics revealed five regularly occurring hemocyte populations. Across individual *P. americana*, the small, non-complex population of cells was the most abundant, closely followed by large cells. The low-complexity group had a variable occurrence across individuals, ranging from approximately 10% of the total population to completely absent **(Figure 1)**. Due to the nature of cockroach stocks, it is possible that this population is present only occasionally due to an unknown, previous bacterial stimulus, due to injury, could arise during different points of development, or could be an artifact of collection (fat bodies). The general shape of whole hemocyte populations from individuals was also broadly variable, with some similar characteristics shared between individuals **(Figure 1A)**. While size and complexity of hemocytes was informative, the variability among individuals contributed to difficulty in using these metrics for cell sorting. While these findings are still novel and informative, they also highlighted the need for alternative, non-species-specific reagents to investigate hemocyte subsets.

Lectins have previously been used to distinguish cell types in other invertebrate systems, including for example, sea urchins^8^, silkworms^12^, and at least once before in other cockroach species^22^. When double staining with WGA and LEA lectins, we observed primarily double positive (+/+) and double negative (-/-) populations, and only sparce single positive (+/- or -/+) populations **(Figure 2A)**. WGA binds both 1,3-N-acetylglucosamine and N-acetylneuraminic acid, while LEA binds only 1,3-N-acetylglucosamine. For this reason, hemocytes that bound LEA would also bind WGA, resulting in a loss of hemocyte population resolution when using both lectins together. It was therefore more informative to evaluate hemocyte subsets based on single stain parameters. Previously, WGA has been described as a general hemocyte marker in *Anopheles gambiae* by Kwon et al. 2021^*11*^. We observed in *P. americana* that there are two hemocyte groups (WGA positive and WGA negative) distinguishable through WGA staining **(Figure 2A)**. It is possible that the WGA negative group were not hemocytes and are instead an artifact of collection, such as fat bodies, as described in some other studies collecting hemocytes through similar methods^23,24^. As no function has currently been attributed to this group, further studies are necessary to determine the identity of these cells. Evaluating hemocytes based on LEA binding capabilities and complexity, we observed three distinct populations with LEA-low non-complex, LEA-low mid-complex, and LEA-high groups **(Figure 2B)**. LEA-high hemocytes were the most abundant, followed by LEA-low non- and mid-complex populations **(Figure 2C)**. While sorting based on these populations does not provide pure hemocyte groups based on morphological differences, LEA-high was enriched in cells with a high cytoplasm to nucleus ratio, LEA-low mid-complex had ovoid “football” shaped cells, and LEA-low non-complex had small cells with a small cytoplasm to nucleus ratio **(Figure 2D)**. These morphologies are consistent with descriptions of granulocytes, plasmatocytes, and prohemocytes respectively as described in other insects^25-27^. It has been previously described in silkworms that granulocytes and plasmatocytes have a high affinity for LEA, and our findings in *P. americana* are consistent with this^12^.

To connect LEA binding to cell function, we measured lysosomal content, ROS production, melanization, and phagocytosis versus LEA binding. Evaluating hemocyte subsets based on additional functional parameters through flow cytometry, we also looked at lysosomal content and ROS production. LysoTracker™ is membrane permeable and fluoresces in acidic environments, acting as a proxy for lysosomal content of cells. *Periplaneta americana* hemocytes had a distinct population of LysoTracker™ high cells **(Figure 3A)**, making up between 14 and 33 percent of the total hemocyte population **(Figure 3D)**. Predictably, when evaluating whole hemocyte populations with LysoTracker™, we found that the more granular hemocytes had the greatest lysosomal content **(Figure 3B)**, while in contrast, the LysoTracker™ negative cells were relatively simple cells **(Figure 3C)**. While expected, this metric provided a means of identifying two distinct hemocyte populations, lyso-high and lyso-low, based on potential cell function. Generally, the high lysosomal content of hemocyte groups aligns with descriptions of granulocytes^29^, plasmatocytes^30^, and oenocytoids.

DCFDA is a membrane permeable ROS indicator, primarily for H_2_O_2,_ in addition to some other reactive oxygen species^28^. There were three distinct populations with varying ROS levels based on DCFDA staining **(Figure 4A)**. The ROS low group was the most abundant, and likely has base ROS levels from metabolic processes. ROS-high granular and non-complex groups are approximately equal in relative abundance, with greater variability in the ROS-high group **(Figure 4B)**. While there was variability in fluorescent intensity, the ROS-high granular group had a higher mean fluorescence than the non-complex group **(Figure 4C)**. The respective roles of hemocytes subsets in ROS production is generally not well understood; however, it has recently been described in *Drosophila* that prohemocytes and crystal cells had both rapid and strong ROS production^31^. This description of ROS production from prohemocytes, which typically are small and non-complex, is consistent with our observations in *P. americana* with our ROS-high low complexity group. While *P. americana* have not been reported to have crystal cells, high ROS production and high granularity would also be consistent with granulocyte and oenocytoid cell types. Additionally, we saw that complex and simple ROS-producing cells also had distinct LEA binding capabilities **(Figure 4D)**, with an LEA-low and LEA-high ROS producing group. These findings together suggest that *P. americana* hemocytes may be separated into a prohemocyte-like and granulocyte-like population based on ROS production and LEA binding capacity.

In other insects, oenocytoids, lamellocytes, and crystal cells have been reported as the prophenoloxidase-producing hemocytes. These enzymes are released after recognition of a PAMP, and induce a proteolytic cascade, leading to the production of cytotoxic melanin^32,33^. *In vivo*, melanization is an immunological process that is commonly paired with nodulation and encapsulation, which also involves the killing and walling off pathogens^34^. From a physiological perspective, oenocytoids or crystal cells would lyse following PAMP recognition, releasing prophenoloxidase into the extracellular matrix. In our assay system, ethanol serves a dual purpose of irreversibly activating prophenoloxidase and fixing the cell membrane to prevent lysis. Additionally, providing L-DOPA (the substrate) allows for melanization inside of cells producing prophenoloxidase, effectively staining melanizing cells without the need for cell lysis. As such, a darkening of the cytoplasm in our sorted cell populations serves as a functional readout of phenoloxidase expression. Our findings demonstrate that the cell population with the highest level of LEA binding was enriched in the most cells containing prophenoloxidase **(Figure 6A)**. In addition to this enrichment in prophenoloxidase-expressing cells, this subset also contained more clustered cells than other subsets, which might be indicative of nodule-forming activity **(Figure 6B)**. It is also worth noting that LEA-high hemocytes are a large group, which could be split into two subsets, the more complex and non-complex. It is possible that this subset is more heterogeneous than initially indicated based on this gating strategy. Thus, more tools are needed to distinguish cells in this subset as well. More experimental evidence is necessary to connect this *in vitro* cluster formation with nodulation or encapsulation. To our knowledge, ours is the first study to connect melanization to hemocyte lectin binding capacity; however, the morphology of the cells expressing prophenoloxidase in our study is consistent with the previous description of oenocytoids^35,36^, or functionally crystal cells (containing prophenoloxidase)^17^, in other systems.

In mammalian systems and some insect species, phagocytosis occurs within an hour of exposure to stimuli^37,38^. To our surprise, in some initial assays, we did not observe phagocytosis in the *P. americana* within this span of time. In some insects, phagocytosis this has been documented to span a much longer window of time^39^. To measure phagocytosis in *P. americana*, we used a time course spanning 24 hours using *S. aureus* conjugated pHrodo™ BioParticles®. This approach allowed us to monitor both phagocytosis and acidification, as the particles are only fluorescently detectible through flow cytometry when in an acidic environment, such as the phago-lysosome. In our studies, phagocytosis first became detectable at approximately 2 hours, and progressively increased over the course of 24 hours. Generally, the majority of particle uptake occurred in the hemocyte population with high LEA-binding capabilities. However, at later time points, there is a second population with lower LEA-binding capabilities phagocytosing particles. These findings indicate that there is a primary, or more active, phagocyte distinguished by phagocytic capabilities and high LEA binding capacity; however, there may be a secondary subset of hemocytes that are phagocytic to a lesser or delayed degree. The kinetics of phagocytosis in *P. americana* investigated through flow cytometry are consistent with phagocytosis kinetics found in some other insects^39^. To our knowledge, we are the first to use LEA as a cell marker for phagocytic hemocytes in insects. Previously, LEA binding has been associated with phagocytic immune cells in oysters^40^, but was not found in shrimp^41^,indicating that lectin binding and phagocytic capabilities of immune cells are variable across invertebrate species. It would therefore be valuable to evaluate lectin binding in association with functional aspects across multiple insect species.

## 5. Conclusions and Future Directions

Our results indicate that there are at least three distinct subsets of hemocytes that can be distinguished through flow cytometry and lectin binding capabilities in *P. americana*. Additionally, we attribute some functionality to these subsets, with our data indicating that granular hemocytes and hemocytes with high affinity for LEA are enriched in cells capable of phagocytosis, melanization, and production of reactive oxygen species. We hypothesize that the non-complex group found in our studies are prohemocytes; however, further functional and transcriptional analysis would be valuable to more precisely determine the identities and functions of these cells. Collectively, our studies provide a reproducible method for investigation of hemocyte and immunological functions in *P. americana* through flow cytometry. These methods are accessible, cost-effective, and may be applied across multiple insect species. Additionally, our data support lectins and functional indicators as potential cell markers for hemocyte populations in systems where antibodies and genomic studies may not be available or practical. In the future, additional functional assays, infection assays, and gene expression studies are necessary to verify the identity and function of hemocytes in *P. americana*.

## Acknowledgements

The authors thank Marla Eva for essential animal care and training, and the AUCVM Flow Cytometry Facility (RRID:SCR_025507) for assistance in cell sorting and analysis. We also thank the National Science Foundation (NSF-IOS-2123655-BIO) and the USDA (AgrSEED 066470972) for funding this research.

## Notes

### Competing Interest Statement

The authors have declared no competing interest.

