## Supplemental figures 1 and 2 for "A flow cytometric approach to identifying the relative abundance and functional capacities of hemocyte subsets in the American cockroach, *Periplaneta americana* (L.)"

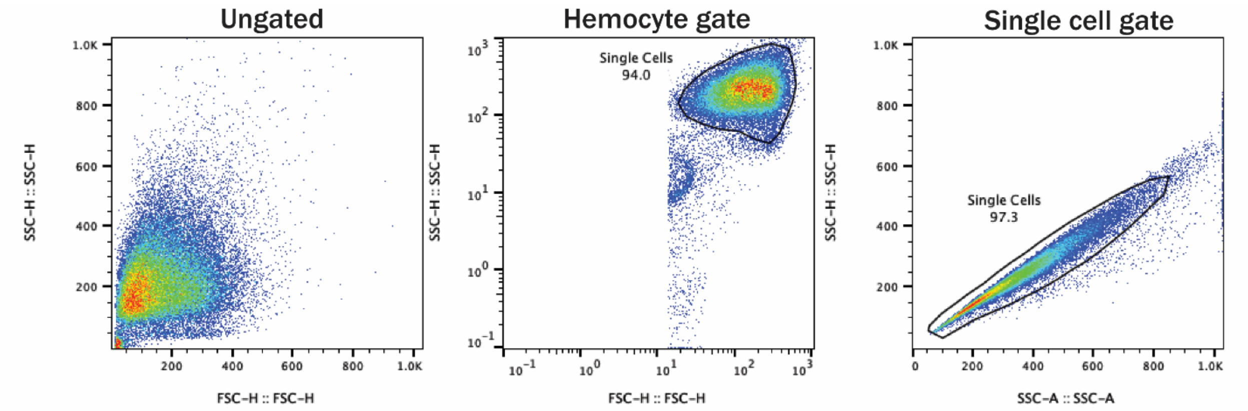


**Supplemental Figure 1.** Example of gating strategy prior to data analysis. The hemocyte gate was separated from debris on a log scale of forward and side scatter. Single cells were determined by viewing hemocytes on a side scatter height and side scatter area graph. Hemocytes outside of the primary cluster were identified as doublets and excluded from analysis. Gates were automatically drawn using the FlowJo 10 auto-gate feature.

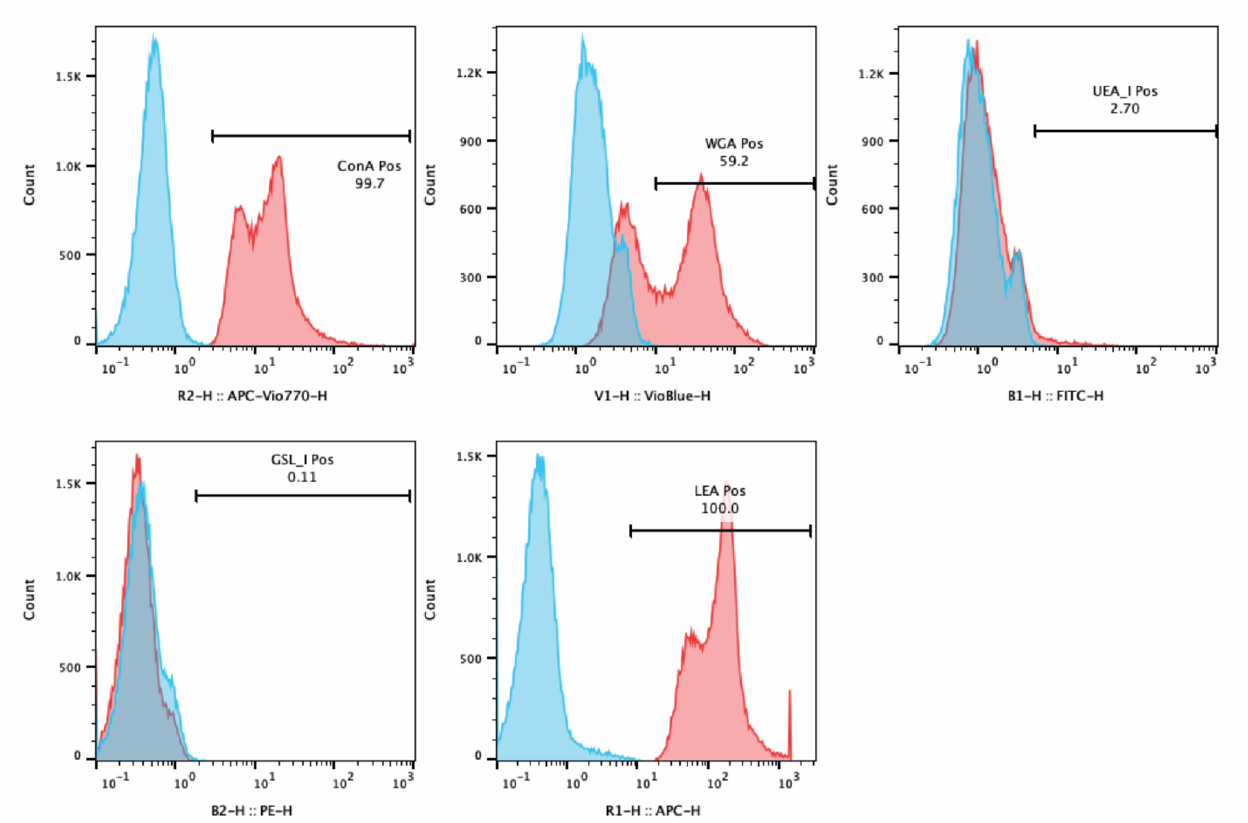


**Supplemental Figure 2.** Lectin binding capabilities of hemocytes. Unstained samples are in blue, and samples incubated with lectins are red. Hemocytes bound ConA, WGA, and LEA. UEA I and GSL I binding was not observed.
